# Exploring the Mechanism of Gentiana in Treating Pancreatic Cancer Based on Network Pharmacology and Molecular Docking Techniques

**DOI:** 10.1101/2024.07.18.604197

**Authors:** Yuanyuan Qian, Zhaojunli Wang, Jiancheng Ji

## Abstract

**Objective:** This study aims to investigate the mechanism of Gentiana in treating pancreatic cancer using network pharmacology and molecular docking techniques.

**Methods:** Active compounds of Gentiana were screened from the Traditional Chinese Medicine Systems Pharmacology (TCMSP) database. The 3D structures of the active compounds were downloaded from the PubChem database. Reverse docking was performed using the PharmMapper database to identify potential target proteins. Differential gene expression data related to colorectal cancer were obtained from the Gene Expression Omnibus (GEO) database, and differentially expressed genes were selected. Venn diagram analysis was employed to identify common genes between the protein targets and differentially expressed genes. Gene Ontology (GO) and Kyoto Encyclopedia of Genes and Genomes (KEGG) analyses were conducted using the Database for Annotation, Visualization, and Integrated Discovery (DAVID) tool. Molecular docking was performed using ChemDraw 20.0, AutoDock, and PyMOL.

**Results:** A total of 72 common genes and 15 signaling pathways were identified from the reverse docking data of Gentiana and the pancreatic cancer dataset (GSE196009). Molecular docking results demonstrated favorable binding energies between the active compounds of Gentiana and proteins 1og5, 1pq2, 2bxr, 2bk3, 1u3w, 1wma, 1wuu, 1tdi, 1mlw, 1egc, 1s1p, 1f12, 1m51, 1kqu, 1ls6, 1ry0, 1nhx, and 1db4.

**Conclusion:** Gentiana may exert its therapeutic effects on pancreatic cancer through a multi-component, multi-target, and multi-pathway mechanism.

## Introduction

Pancreatic cancer, known as the “king of cancers” in the field of oncology, is one of the common malignant tumors in the digestive system[1]. It presents with obscure and atypical clinical symptoms, making it a challenging malignancy to diagnose and treat in the digestive tract. Approximately 90% of pancreatic cancers originate from the ductal epithelium of the exocrine pancreas[2]. Its incidence and mortality rates have been significantly increasing in recent years [3]. Its incidence and mortality rates have been significantly increasing in recent years [4]. It predominantly affects middle-aged and elderly individuals[5], with a higher prevalence in males than premenopausal females, but similar incidence rates in postmenopausal females[6–8].

Gentiana, a perennial herb belonging to the Gentianaceae family, features white-yellow roots that resemble ropes and upright stems with a purple-brown hue. Its leaves grow in pairs, and they are ovate or lanceolate with rough edges and veins on the underside. The flowers cluster at the stem tips or leaf axils, and the bracts are lanceolate, nearly equal in length to the calyx, which is bell-shaped. It is found in meadows, thickets, or forest edges [9, 10]. Traditionally, Gentiana is used to treat conditions related to liver heat, epilepsy, mania, encephalitis B, headache, red eyes, sore throat, jaundice, dysentery, abscesses, and swelling and pain in the scrotum and vulva[9, 11, 12]. Research has revealed that extracts of Gentiana root possess new pharmacological activities, exhibiting anti-proliferative effects on tumor cells in both cultured cells and animal models[13].

Network pharmacology is a novel discipline based on systems biology, which employs network analysis of biological systems to select specific signal nodes for the design of multi-target drug molecules[14]. Through the multi-pathway regulation of signaling pathways, network pharmacology aims to enhance the therapeutic effects of drugs while reducing their toxic side effects, thereby increasing the success rate of new drug clinical trials and lowering drug development costs[15]. With the rise of network pharmacology, its overall systemic characteristics align with the holistic view and the principles of pattern identification and treatment in traditional Chinese medicine (TCM). It has been widely applied in TCM research, providing new insights into the study of complex herbal systems and offering new technological support for rational drug use and new drug development [16, 17].

Molecular docking is a theoretical simulation method that studies the interaction between molecules by investigating the characteristics of the receptor and the interaction mode and affinity between the receptor and the drug molecule[18, 19]. It primarily focuses on the interaction between molecules and predicts their binding modes and affinities[20].

This study aims to explore the potential therapeutic targets of Gentiana in treating pancreatic cancer through the joint analysis of network pharmacology and gene chip data. Additionally, molecular docking techniques will be employed to investigate the binding capacity between the active compounds of Gentiana and target proteins, validating the potential of Gentiana in treating pancreatic cancer.

## 1 Materials and Methods

### 1.1 Screening of Active Ingredients in Gentiana

The active ingredients of Gentiana were screened from the TCMSP database (http://tcmspw.com/tcmsp.php) based on the criteria of oral bioavailability (OB) ≥ 30.00 and drug-likeness (DL) ≥ 0.18.

### 1.2 Screening of Target Proteins for Active Ingredients

The 3D structures of the active ingredients of Gentiana were downloaded from the PubChem database (https://pubchem.ncbi.nlm.nih.gov/). The file format was converted to mol2 format using openBabel software. The active ingredients were then uploaded to the PharmMapper database (http://www.lilab-ecust.cn/pharmmapper/) for reverse docking. Target proteins were selected and merged based on a z-score > 1. The Gentiana - Active Ingredients - Target Proteins (PDB ID) network diagram was drawn using Cytoscape.

### 1.3 Conversion of PDB IDs to SYMBOL IDs

PDB IDs were converted to SYMBOL IDs using the Uniprot database (https://www.uniprot.org/) and the DAVID database (https://david.ncifcrf.gov/home.jsp).

### 1.4 Screening and Processing of Pancreatic Cancer Data

Pancreatic cancer datasets containing normal and cancerous tissues were selected from the GEO database (https://www.ncbi.nlm.nih.gov/geo/). Differential gene expression analysis was performed to identify differentially expressed genes. Genes with |Log(FC)| > 1 and P-value < 0.05 were selected, and a volcano plot was generated.

### 1.5 Selection of Common Genes

The Venn diagram was used to select common genes from steps 1.3 and 1.4, and the Gentiana - Active Ingredients - Target Proteins (PDB ID) - Gene network diagram was drawn using Cytoscape.

### 1.6 GO and KEGG Analysis

The common genes selected were subjected to GO and KEGG analysis using the DAVID database, and genes with a P-value < 0.05 were filtered. Biological Process (BP) and KEGG pathway analyses were conducted, and the results were visualized using Microbiux (http://www.bioinformatics.com.cn/). A network diagram of Gentiana - Active Ingredients - Target Proteins (PDB ID) - Genes - Pathways was drawn using Cytoscape.

### 1.7 Molecular Docking

1. Protein and Ligand Download: The 2D structures of the active ingredients of Gentiana were downloaded from the PubChem database. The 3D structures of target proteins were obtained from the Protein Data Bank (PDB) (https://www.rcsb.org/) and the AlphaFold protein prediction database (https://alphafold.com/).
2. Preprocessing: The active ingredients of Gentiana were imported into ChemDraw 20.0 software for energy minimization simulation. The target protein files were imported into PyMOL software for desalting, hydrogenation, and removal of ligands. Multimeric proteins were split into monomers.
3. Molecular Docking: The target proteins were processed for hydrogenation and charging using AutoDockTools. The active ingredients of Gentiana were processed for hydrogenation, charging, and root addition. Molecular docking was performed using autogrid4 and autodock4.
4. Visualization of Docking Results: The docking result files were converted to PDB format using openBabel software, and visualization was performed using PyMOL software.

## 2 Results

### 2.1 Screening of Active Components and Target Proteins of Gentiana

The active components of Gentiana were screened from the TCMSP database, resulting in 10 active components based on the screening criteria of OB ≥ 30.00 and DL ≥ 0.18. The 3D structures of these active components were downloaded from the PubChem database and subjected to reverse docking using the PharmMapper database. A total of 360 target proteins were screened with a selection criterion of z-score > 1 (Figure 1). The PDB IDs were then converted to SYMBOL IDs using the Uniprot and DAVID databases, resulting in 263 SYMBOL IDs, five of which were not found in the Uniprot database for humans.

**Figure 1:**
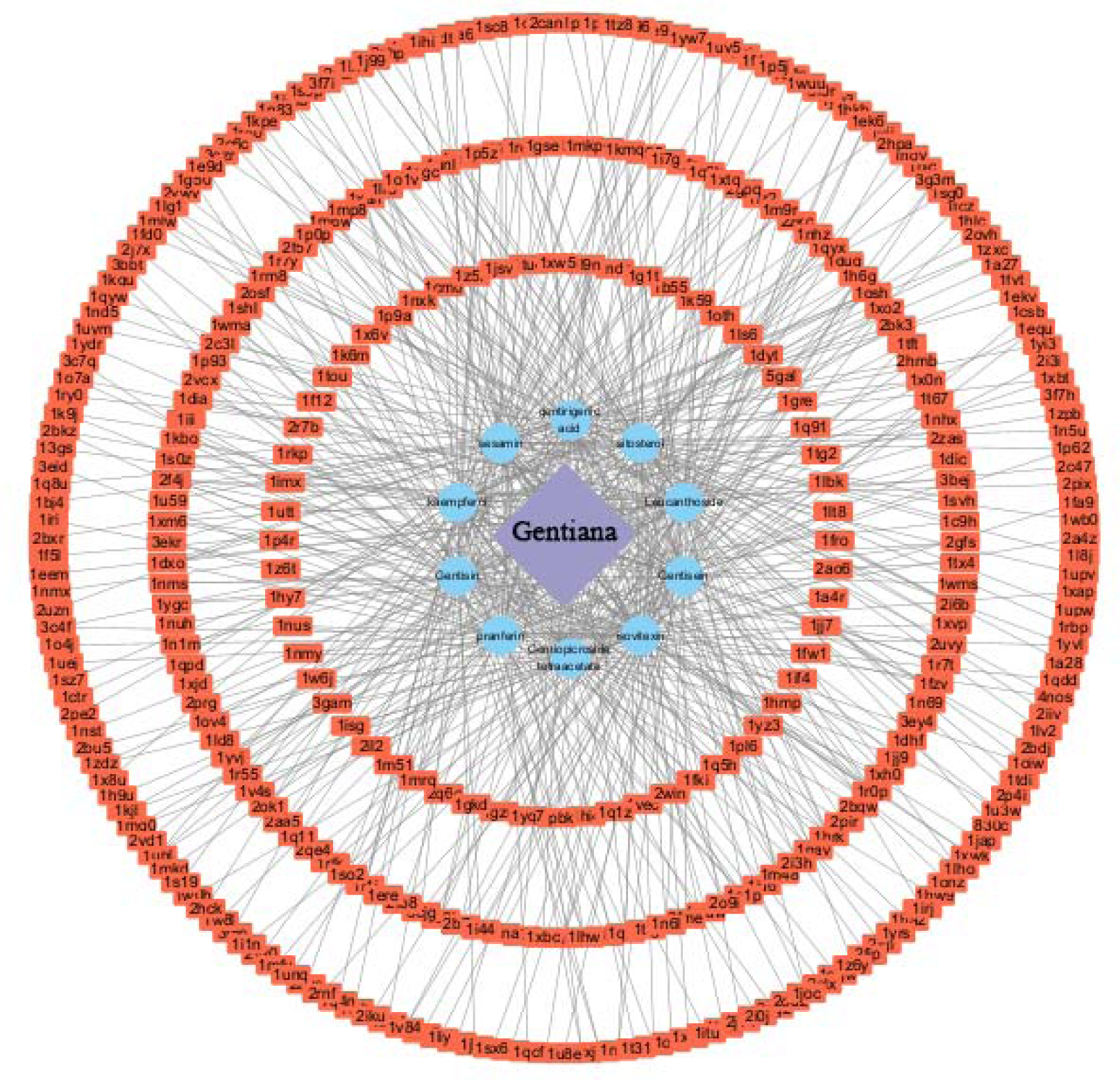
Network diagram of Gentiana - Active ingredients - Target proteins (PDB ID): Visualized using Cytoscape software to illustrate the interaction relationships between Gentiana, its active ingredients, and target proteins.

### 2.2 Analysis of Pancreatic Cancer Dataset

The GSE196009 gene microarray dataset was screened from the GEO database, and differential gene analysis was performed. Based on the selection criteria of |Log(FC)| > 1 and P-value < 0.05, 5050 differentially expressed genes were identified (2584 upregulated genes and 2466 downregulated genes) (Figure 2).

**Figure 2:**
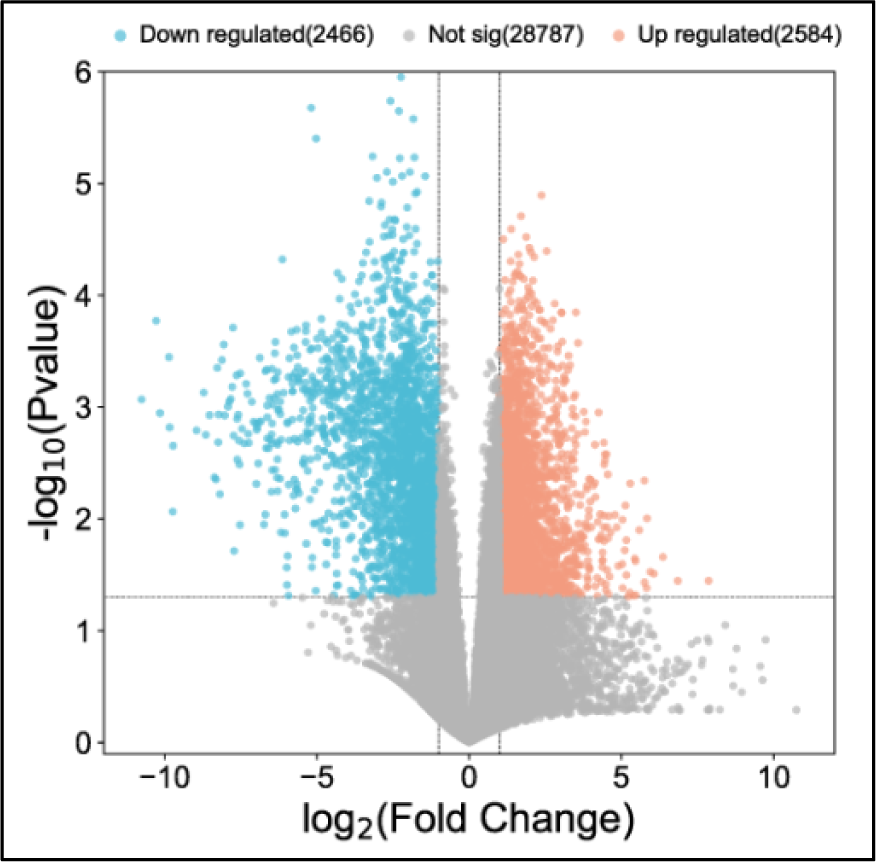
Volcano plot of gene expression in pancreatic cancer GSE196009 gene chip dataset.

### 2.3 Analysis of Common Genes

The Venn diagram was used to screen common genes between the target genes of Gentiana active components and the differential genes in the GSE196009 gene microarray of pancreatic cancer, identifying 72 different genes (Figure 3A). The Gentiana - Active Components - Target Protein - Common Gene network was constructed using Cytoscape software (Figure 3B).

**Figure 3:**
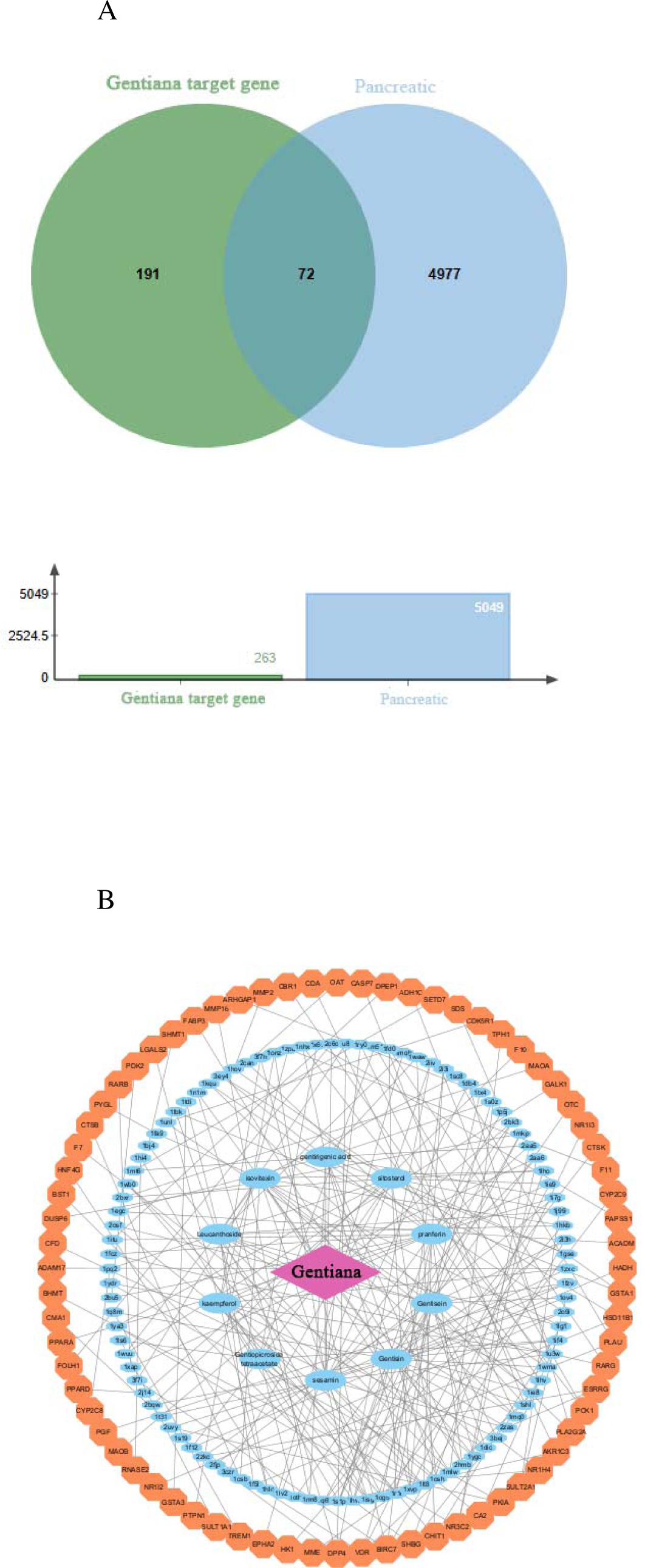
Network relationship diagram between Gentiana and pancreatic cancer target genes. A: Venn diagram showing the common genes between Gentiana active ingredients and differentially expressed genes in the pancreatic cancer GSE196009 gene chip dataset; B: Network diagram of Gentiana - Active ingredients - Target proteins - Common genes.

### 2.4 GO and KEGG Analysis

The selected common genes were imported into the DAVID database for GO and KEGG analysis (Figure 4A, Figure 4B). The Gentiana - Active Components - Target Protein (PDB ID) - Gene - Pathway network was drawn using Cytoscape (Figure 4C).

**Figure 4:**
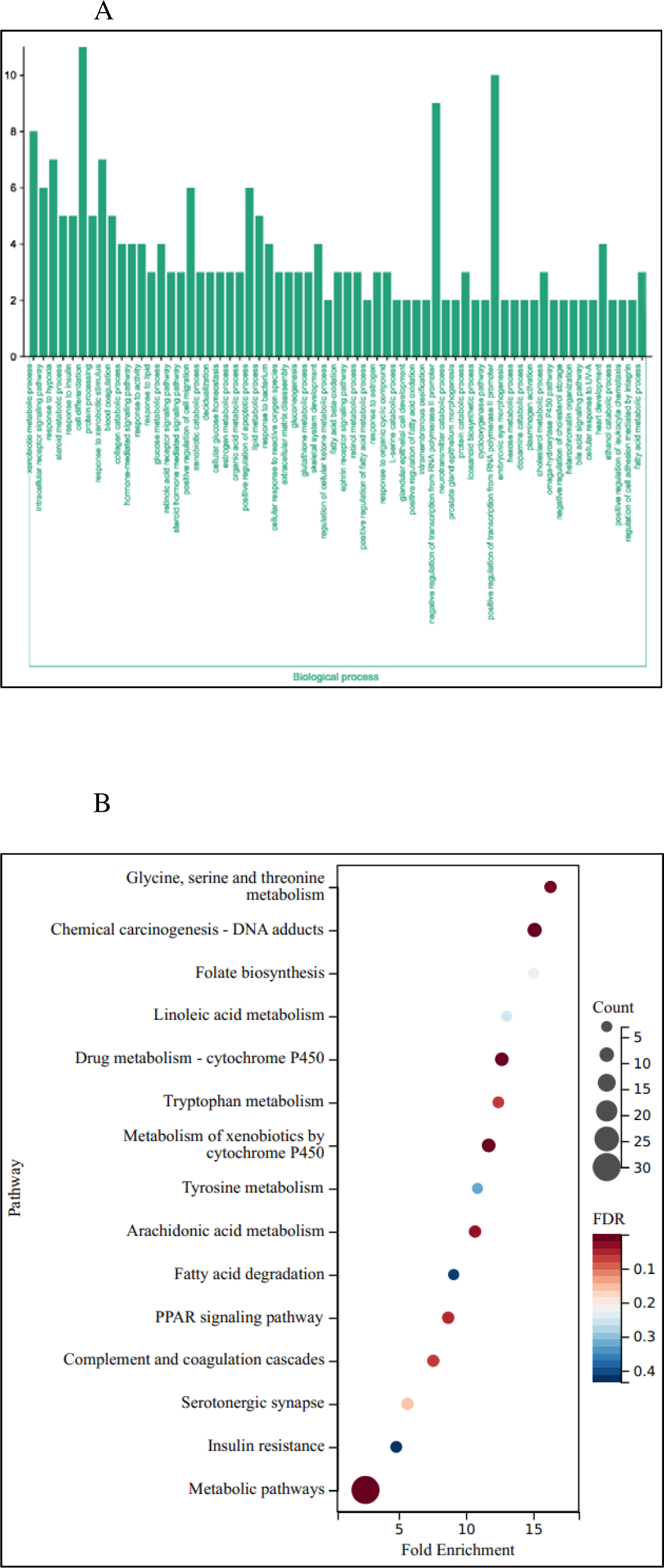
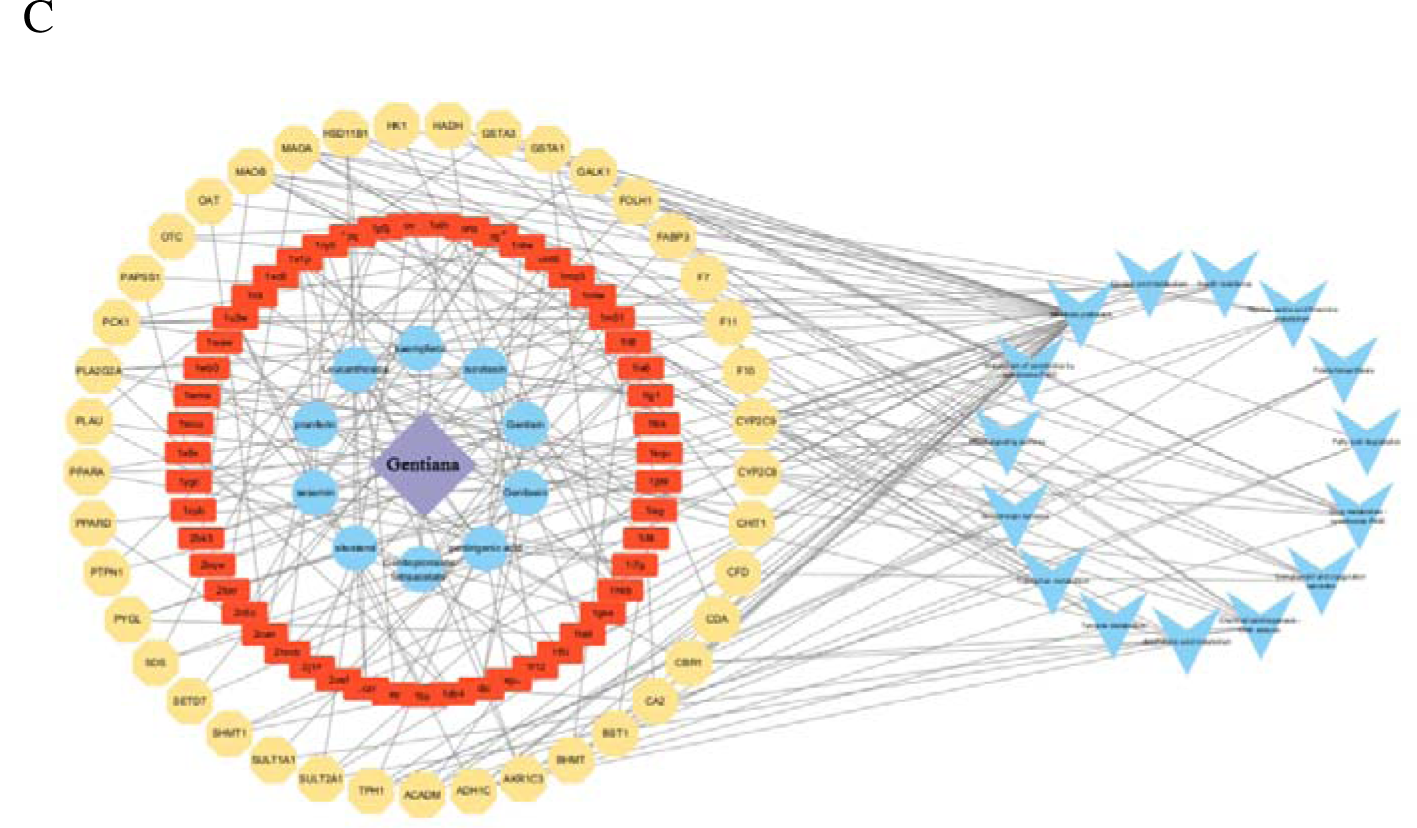
Network relationship diagram between Gentiana, pancreatic cancer target genes, and signaling pathways. A: Visualization of GO (Gene Ontology) Biological Process results; B: Visualization of KEGG (Kyoto Encyclopedia of Genes and Genomes) pathway analysis results; C: Network diagram of Gentiana - Active ingredients - Proteins - Common genes - Target proteins.

### 2.5 Molecular Docking

Eighteen target genes were selected based on the KEGG results and the 18 corresponding target proteins were selected based on the results of section 3.1. Proteins and active components of Gentiana were preprocessed using ChemDraw 20.0 and PyMOL software, respectively. Molecular docking was performed using AutoDock software, and visualization was performed using PyMOL software.

Gentiopicroside tetraacetate formed hydrogen bonds with Glutamic acid (GLU)-110, Isoleucine (ILE)-108, Alanine (ALA)-107, Methionine (MET)-26 in 1f12, and GLN-451, Tryptophan (TRP)-450, LEU-508, LYS-510 in 1m51 (Figure S1).

Gentirigenic acid formed hydrogen bonds with LYS-282, LYS-148, LYS-187 in 1egc, and LYS-304, Glycine (GLY)-302, Proline (PRO)-298 in 1nhx (Figure S2). Gentisein formed hydrogen bonds with ALA-218, LEU-268, Aspartic acid (ASP)-50, GLN-222 in 1s1p (Figure S3). Gentisin formed hydrogen bonds with Serine (SER)-217, Tyrosine (TYR)-55, GLN-222, LYS-270 in 1s1p; Cysteine (CYS)-172, TYR-435 in 2bk3; and Valine (VAL)-230, Asparagine (ASN)-89, TYR-193 in 1wma (Figure S4). Isovitexin formed hydrogen bonds with Threonine (THR)-265, GLU-317, ALA-309 in 1mlw; Histidine (HIS)-88, GLN-87 in 1wuu; and Phenylalanine (PHE)-94, GLN-105 in 1wma (Figure S5). Kaempferol formed hydrogen bonds with SER-221, LYS-270, LEU-268 in 1s1p; LEU-116, VAL-292 in 1u3w; and PHE-100, GLY-98 in 1og5 (Figure S6). Leucanthoside formed hydrogen bonds with GLU-340 in 1mlw and GLY-14, ARG-13, GLU-169, LEU-107 in 1tdi (Figure S7). Pranferin formed hydrogen bonds with GLN-222, TYR-55, HIS-117 in 1s1p; PHE-100, GLY-98 in 1og5; and HIS-47, ASP-48 in 1kqu (Figure S8). Sesamin forms hydrogen bonds with LYS-52 and GLY-29 in 1db4; LEU-219, SER-217, LEU-268, GLN-222 in 1ry0; GLN-356, LEU-361, VAL-436, ARG-433 in 1og5; ILE-199, ILE-198, GLN-206 in 2bk3; and THR-178, GLY-201, LEU-200 in 1u3w (Figure S9). Sitosterol forms hydrogen bonds with LYS-474 in 1pq2 and THR-205, PHE-208 in 2bxr (Figure S10). The above molecular docking data are in Table 1.

**Table 1.**
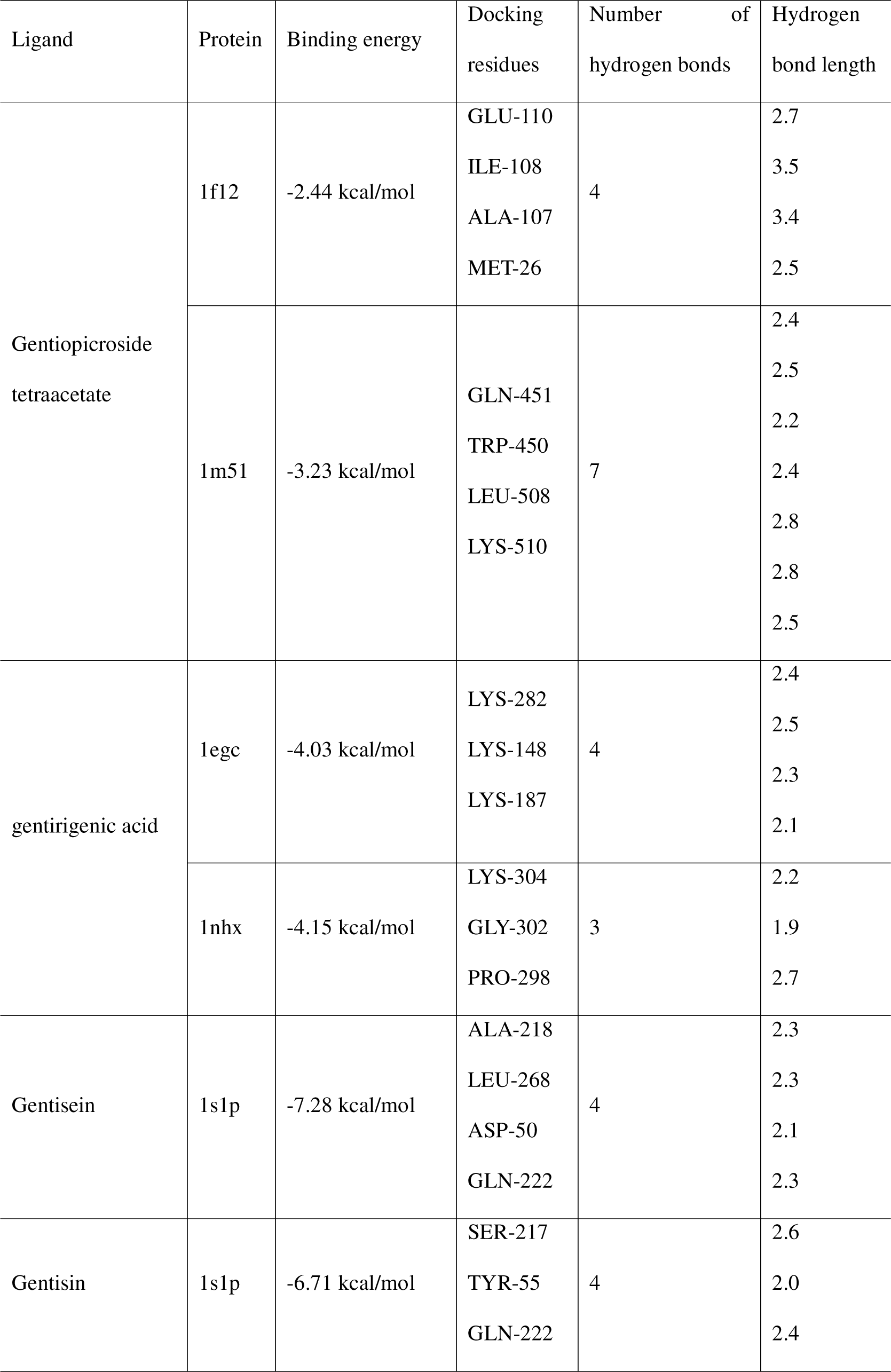
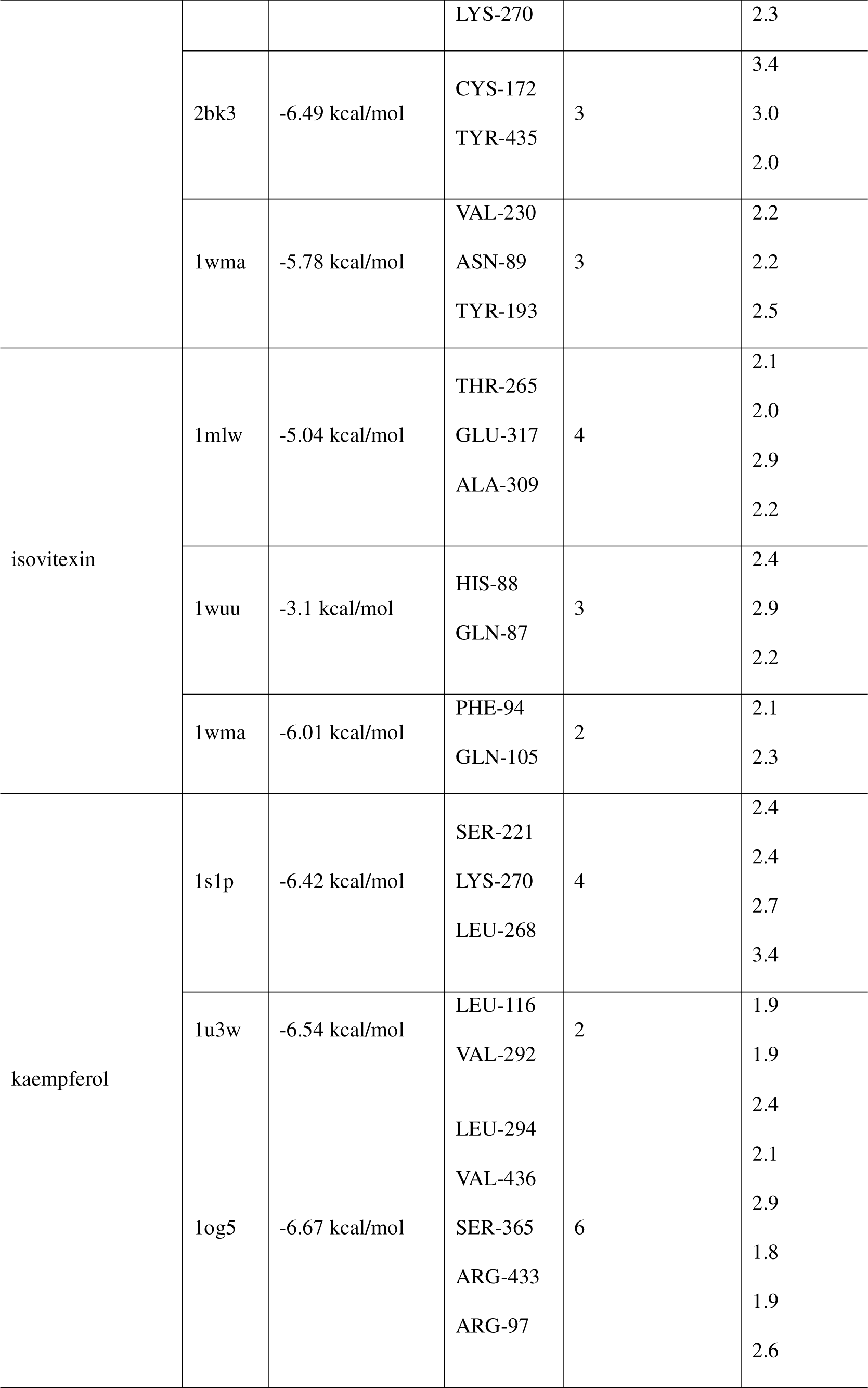
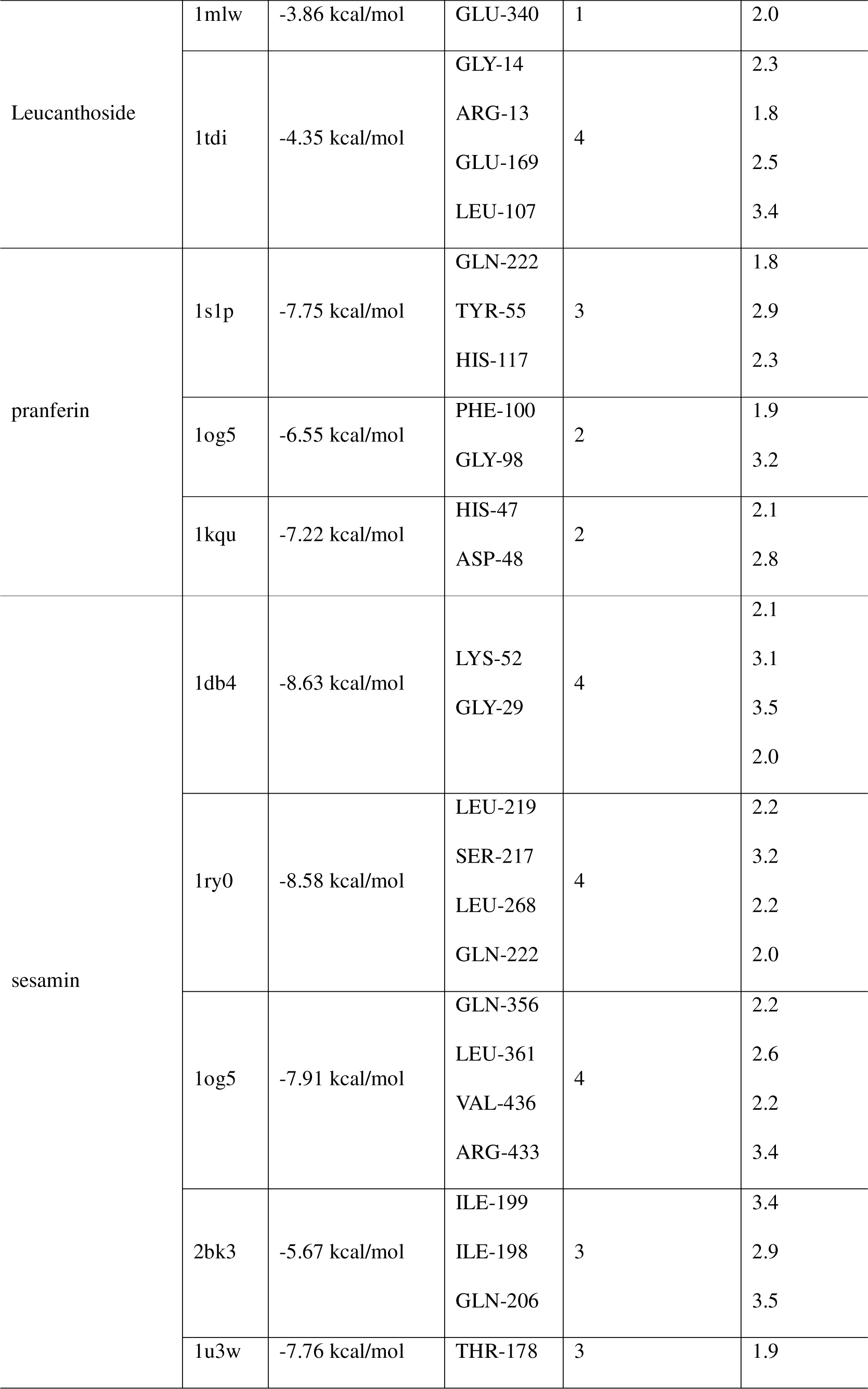
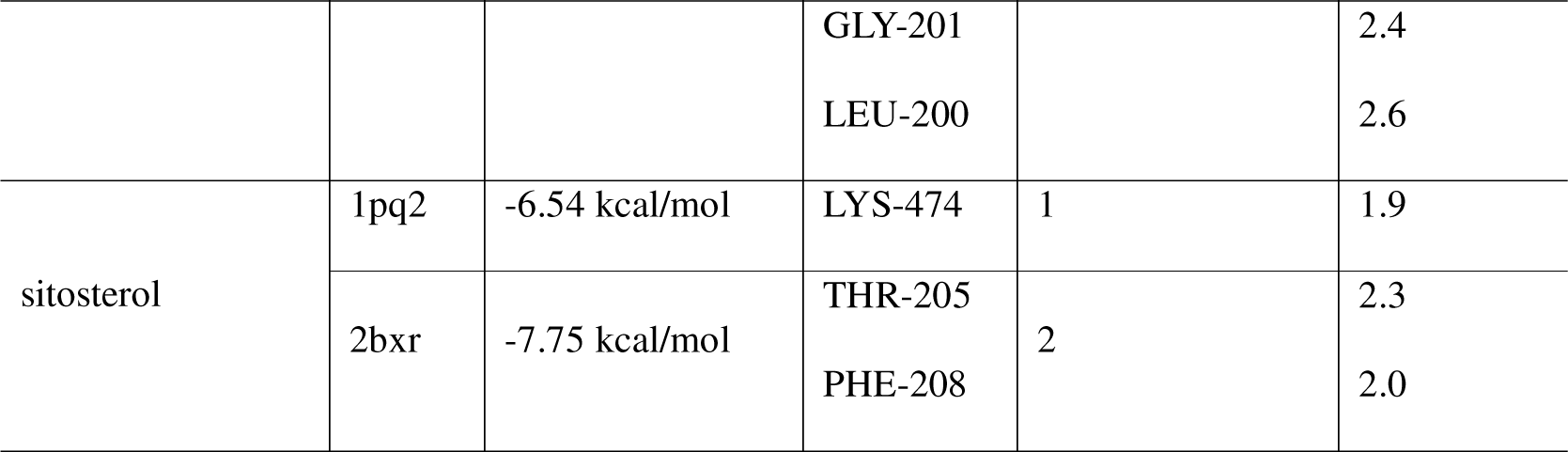
Molecular docking data.

## 3 Discussion

Pancreatic cancer is characterized by its high malignancy, low surgical resection rate, and poor prognosis[2, 21, 22]. Although surgery remains the primary treatment method, the late diagnosis of pancreatic cancer often leads to the loss of curative opportunities, necessitating comprehensive treatment options[23]. To date, like most cancers, there is still no highly effective and universally applicable comprehensive treatment approach. The current comprehensive treatment for pancreatic cancer still relies primarily on surgical intervention with radiotherapy and chemotherapy as adjuvants [22, 23]. Chinese herbal medicine possesses characteristics such as effectiveness, mildness, and minimal side effects, making it suitable for combination therapy with radiation and chemotherapy to improve the quality of life of patients. Experimental studies have demonstrated that active compounds found in Gentiana, such as sesamin, kaempferol, and isovitexin, can effectively inhibit the proliferation, induce apoptosis, and suppress migration of cancer cells[24–27], possibly through the mTORC1 signaling pathway, PI3K/AKT pathway, and Wnt/β-catenin signaling pathway, thereby inhibiting cancer cell activity, proliferation, and migration[25].

In this study, network pharmacology techniques were employed to comprehensively evaluate the relationship between Gentiana and pancreatic cancer and predict the molecular mechanisms underlying the therapeutic effects of Gentiana on pancreatic cancer. Through screening in various databases, ten active compounds from Gentiana and 72 potential target genes were identified.

The KEGG analysis revealed potential pathways through which Gentiana may treat pancreatic cancer:

1. The PPAR signaling pathway is activated by fatty acids and their derivatives and belongs to the ligand-activated receptor superfamily of nuclear hormone receptors. Among them, PPARα is a member of the peroxisome proliferator-activated receptors (PPARs) family and plays a critical role in regulating cell proliferation, apoptosis, and tumorigenesis. Intracellular fatty acid-binding proteins (FABPs) belong to a multigene family and are classified into at least three types: liver-type, intestinal-type, and heart-type. They form 14-15 kDa proteins and are believed to be involved in the uptake, intracellular metabolism, and/or transport of long-chain fatty acids. They may also be responsible for regulating cell growth and proliferation. Gentiana may regulate the low expression levels of ACADM, PCK1, and FABP1 genes to increase the secretion of PPARα, PPARβ/δ, and PPARγ transcription factors, thereby enhancing the expression of downstream factors[28, 29].
2. Target genes related to cancer cell metabolism pathways mainly include metabolic pathways, arachidonic acid metabolism, fatty acid degradation, and glycine, serine, and threonine metabolism. Studies have shown that inhibiting arachidonic acid metabolism can suppress pancreatic cancer cell proliferation. Arachidonic acid can be metabolized through three pathways: cyclooxygenases (COX), lipoxygenases (LOX), and cytochrome P450 (CYP450), resulting in the production of numerous metabolites such as prostaglandins, thromboxanes, leukotrienes, hydroxyeicosatetraenoic acids, epoxyeicosatetraenoic acids, etc. These metabolites play important roles in organ function maintenance, inflammatory responses, drug metabolism, and are involved in the development of various diseases, including cardiovascular diseases and cancer. Gentiana may regulate the expression of genes such as CYP2C9 and CBR1, which influence the expression of arachidonic acid, thereby inhibiting pancreatic cancer cell proliferation[30, 31].

In the GO BP enrichment analysis, target genes were mainly enriched in cell metabolism pathways, signal transduction pathways, and cell differentiation pathways. They also showed regulatory effects on processes such as cell differentiation, positive regulation of apoptosis, and positive regulation of cell migration. Metabolism within the body is interconnected and interrelated, and multiple metabolic pathways in various systems, such as the circulatory and nervous systems, can indirectly influence or intervene in the cancer cell environment.

The molecular docking results demonstrated a strong binding affinity between the active compounds of Gentiana and target proteins, with binding energies of less than 5 kJ/mol, indicating a stable interaction between the active compounds and target proteins. This provides some evidence for the potential therapeutic effects of Gentiana on pancreatic cancer.

In summary, the results of this study suggest that Gentiana may exert its therapeutic effects on pancreatic cancer through the synergistic action of multiple active compounds and target genes, as well as the regulation of various signaling pathways and metabolic pathways. However, further clinical experiments and in-depth research are required to validate and interpret these findings and provide more scientific and reliable evidence for the application of Gentiana in the treatment of pancreatic cancer.

## 4 Conclusion

In conclusion, Gentiana may inhibit or influence the growth and development of pancreatic cancer cells through metabolic pathways, signal transduction pathways, and other signaling pathways. It exhibits the characteristic of multi-component, multi-target, and multi-signaling pathway effects in treating pancreatic cancer, providing a preliminary exploration of the biological pathways underlying the therapeutic effects of Gentiana on pancreatic cancer.

As this study is based on predictive analysis, a more comprehensive analysis of all the components, target points, and signaling pathways of Gentiana would require further investigations. Subsequent experiments at the protein level, cell models, animal models, and clinical trials are needed to validate the effects of Gentiana on pancreatic cancer and to refine the theoretical research of this medicinal herb.

## Conflicts of interest

There are no conflicts of interest.

## Authors’ contributions

Yuanyuan Qian participated in the design of the study and review. Zhaojunli Wang and Jiancheng Ji drafted the manuscript and participated in the data collection and analysis. All authors contributed to the article and approved the submitted version.

## Data availability statement

The datasets used and analyzed during the current study are available from the corresponding author upon reasonable request.

## Acknowledgements

Not applicable.

## Funding

None.

## Notes

### Competing Interest Statement

The authors have declared no competing interest.

### Summary of Updates

author affiliations updated:Department of Research and Development, Jilin Ruiguo Technology Co., Ltd, Changchun, 130000, China.

