## Supplementary Material Figure S for "Exploring the Mechanism of Gentiana in Treating Pancreatic Cancer Based on Network Pharmacology and Molecular Docking Techniques"

Figure S1.

A. Gentiopicroside tetraacetate and 1f12 docking result; B. Gentiopicroside tetraacetate and 1m51 docking result. The specific interactions between Gentiopicroside tetraacetate and the amino acid residues in the target proteins 1f12 and 1m51 are shown in the respective docking result figures. These interactions indicate potential binding and stabilization of Gentiopicroside tetraacetate with the target proteins.

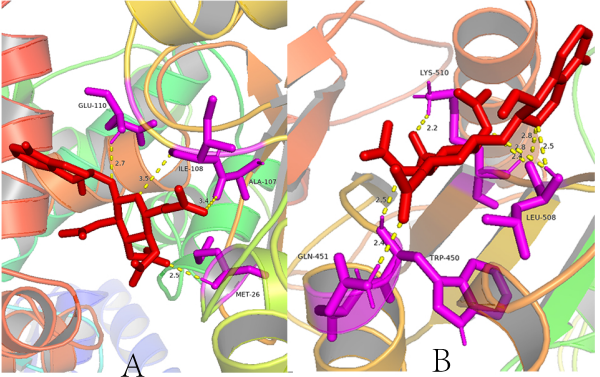

Figure S2.

A. Docking Results of Gentirigenic Acid with 1egc; B. Docking Results of Gentirigenic Acid with 1nhx; The specific details of the binding interactions and binding energies are not provided in the given text. Please refer to the original research for a more comprehensive analysis of the docking results.

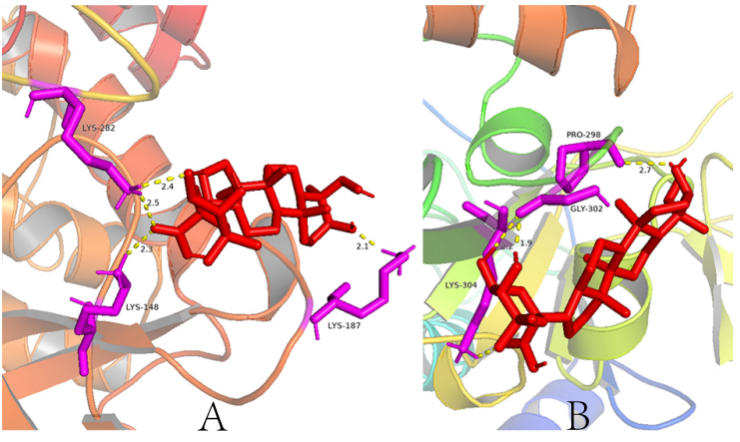

Figure S3. Docking Results of Gentisein with 1s1p.

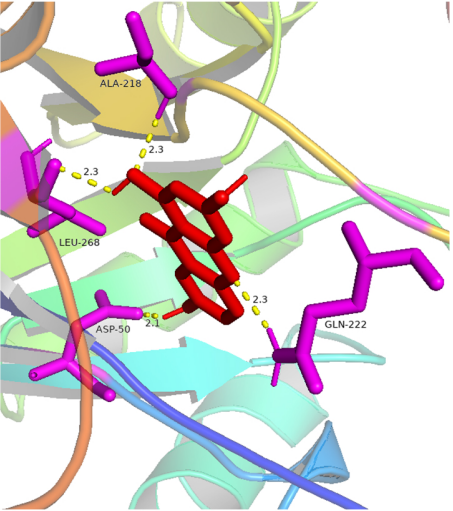

Figure S4. A. Docking Results of Gentisin with 1s1p; B. Docking Results of Gentisin with 1wma; C. Docking Results of Gentisin with 2bk3.

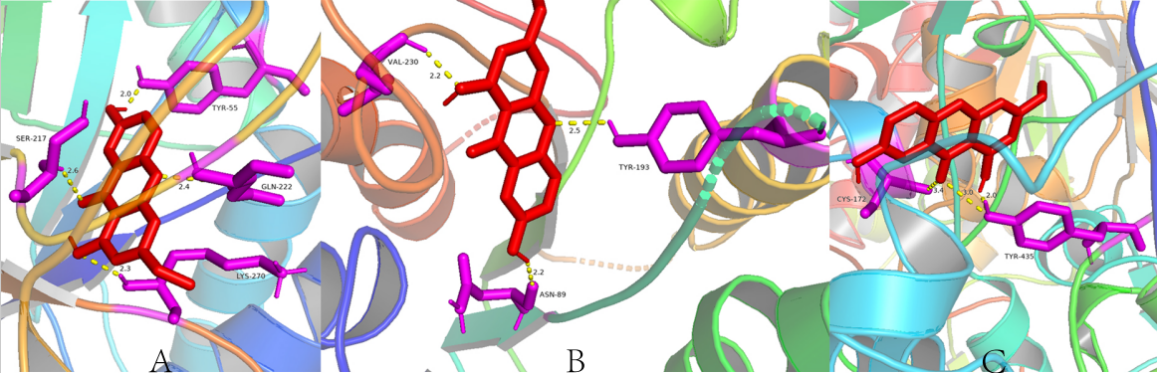

Figure S5. A. Docking Results of Isovitexin with 1mlw; B. Docking Results of Isovitexin with 1wma; C. Docking Results of Isovitexin with 1wuu.

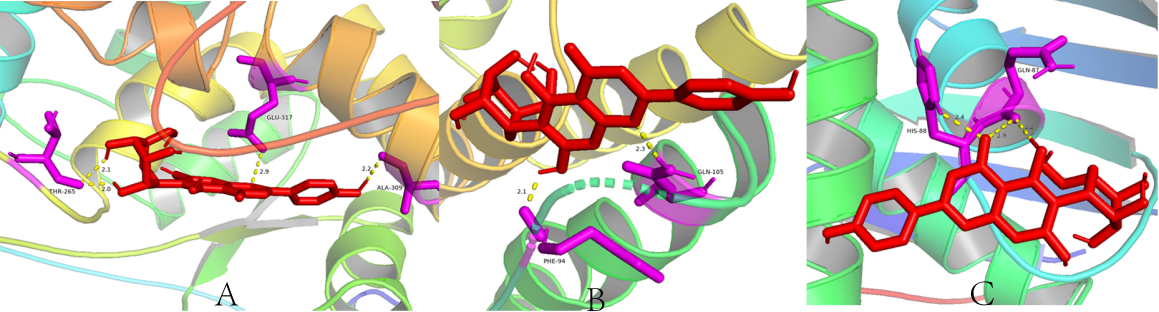

Figure S6. A. Docking Results of Kaempferol with 1og5; B. Docking Results of Kaempferol with 1s1p; C. Docking Results of Kaempferol with 1u3w.

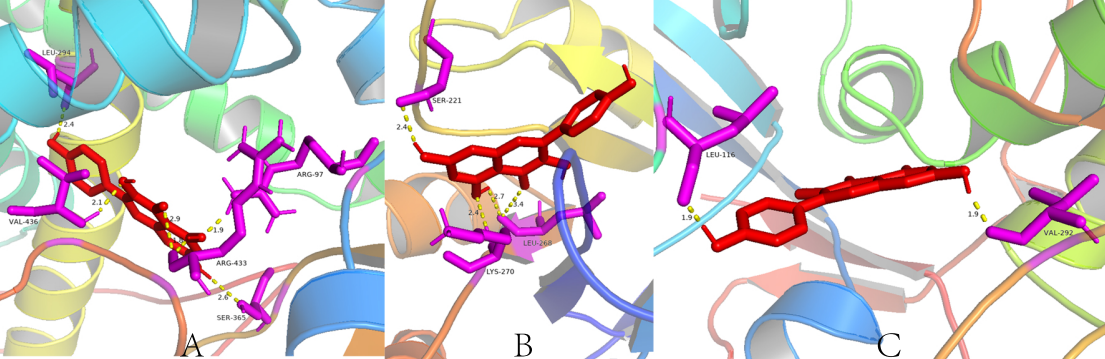

Figure S7. A. Docking Results of Leucanthoside with 1mlw; B. Docking Results of Leucanthoside with 1tdi.

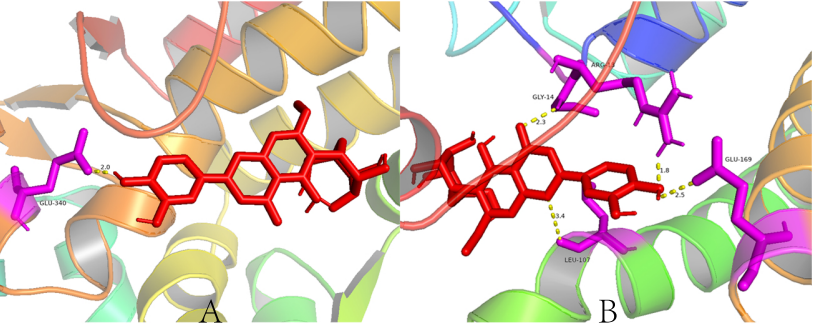

Figure S8. A. Docking Results of Pranferin with 1kqu; B. Docking Results of Pranferin with 1og5; C. Docking Results of Pranferin with 1s1p.

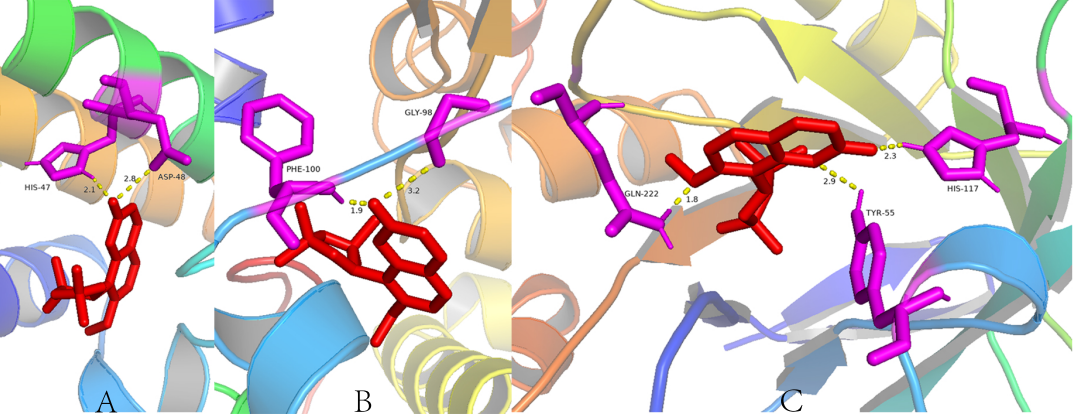

Figure S9. A. Docking Results of Sesamin with 1db4; B. Docking Results of Sesamin with 1og5; C. Docking Results of Sesamin with 1ry0; D. Docking Results of Sesamin with 1u3w; E. Docking Results of Sesamin with 2bk3.

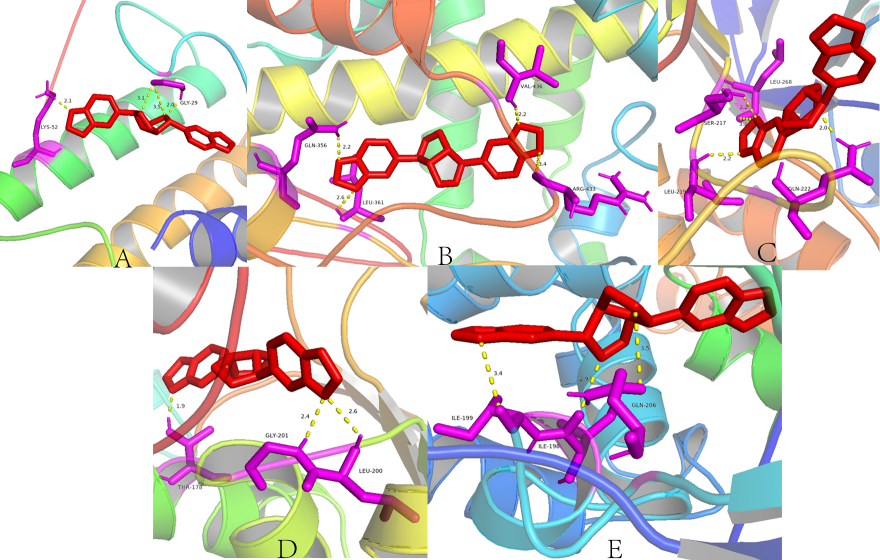

Figure S10. A. Docking Results of Sitosterol with 1pq2; B. Docking Results of Sitosterol with 2bxr.

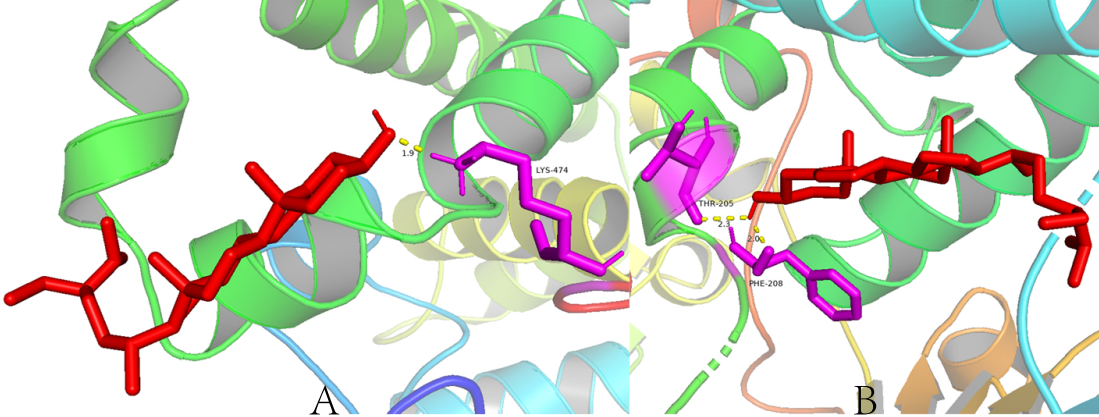
